# Sequencing the orthologs of human autosomal forensic short tandem repeats provides individual- and species-level identification in African great apes

**DOI:** 10.1101/2022.08.03.502616

**Authors:** Ettore Fedele, Jon H. Wetton, Mark A. Jobling

## Abstract

**Background:** Great apes are a global conservation concern, with anthropogenic pressures threatening their survival. Genetic analysis can be used to assess the effects of reduced population sizes and the effectiveness of conservation measures. In humans, autosomal short tandem repeats (aSTRs) are widely used in population genetics and for forensic individual identification and kinship testing. Traditionally, genotyping is length-based via capillary electrophoresis (CE), but there is an increasing move to direct analysis by massively parallel sequencing (MPS). Here we assess in African great ape DNAs the human-based ForenSeq DNA Sequencing Prep Kit, which amplifies multiple loci including 27 aSTRs, prior to sequencing via Illumina technology. We ask whether cross-species genotyping of the orthologs of these loci can provide both individual and (sub)species identification.

**Results:** The Forenseq kit was used to amplify and sequence aSTRs in 52 individuals (14 chimpanzees; 4 bonobos; 16 western lowland, 6 eastern lowland, and 12 mountain gorillas). The orthologs of 24/27 human aSTRs amplified across species, and a core set of thirteen loci could be genotyped in all individuals. Genotypes were individually and (sub)species identifying. Both allelic diversity and the power to discriminate (sub)species were greater when considering STR sequences rather than allele lengths. Comparing human and African great-ape STR sequences with an orangutan outgroup showed general conservation of repeat types and allele size ranges, but variation in repeat array structures and little relationship with the known phylogeny, suggesting stochastic origins of mutations giving rise to diverse imperfect repeat arrays. Interruptions within long repeat arrays in African great apes do not appear to reduce allelic diversity, indicating a possible mutational difference to humans.

**Conclusions:** Despite some variability in amplification success, orthologs of most human aSTRs in the ForenSeq DNA Sequencing Prep Kit can be analysed in African great apes. MPS of the orthologs of human loci provides better resolution for both individual and (sub)species identification in great apes than standard CE-based approaches, and has the further advantage that there is no need to limit the number and size ranges of analysed loci.

## Background

Habitat loss, disease, climate change and hunting are among the main drivers of localised and global extinctions [1]. As species become increasingly restricted to fragmented habitats it is necessary to assess their viability to support effective management decisions. Increasing global awareness has drawn attention towards the preservation of charismatic flagship species [2], among which the African great apes have been a focal interest: most of these species remain critically endangered throughout their home ranges [3] (Figure 1). However, when threat status is measured merely on the basis of species decline and habitat degradation [4], it can neglect the biological and ecological impacts of shifts in population size and distribution [5]. As populations decline and inbreeding intensifies, high rates of homozygosity spread among groups of individuals [6]. In turn, reduced allelic diversity can affect the adaptive ability of the species and potentially lead to the emergence of genetic defects underpinned by recessive alleles [7].

**Figure 1:**
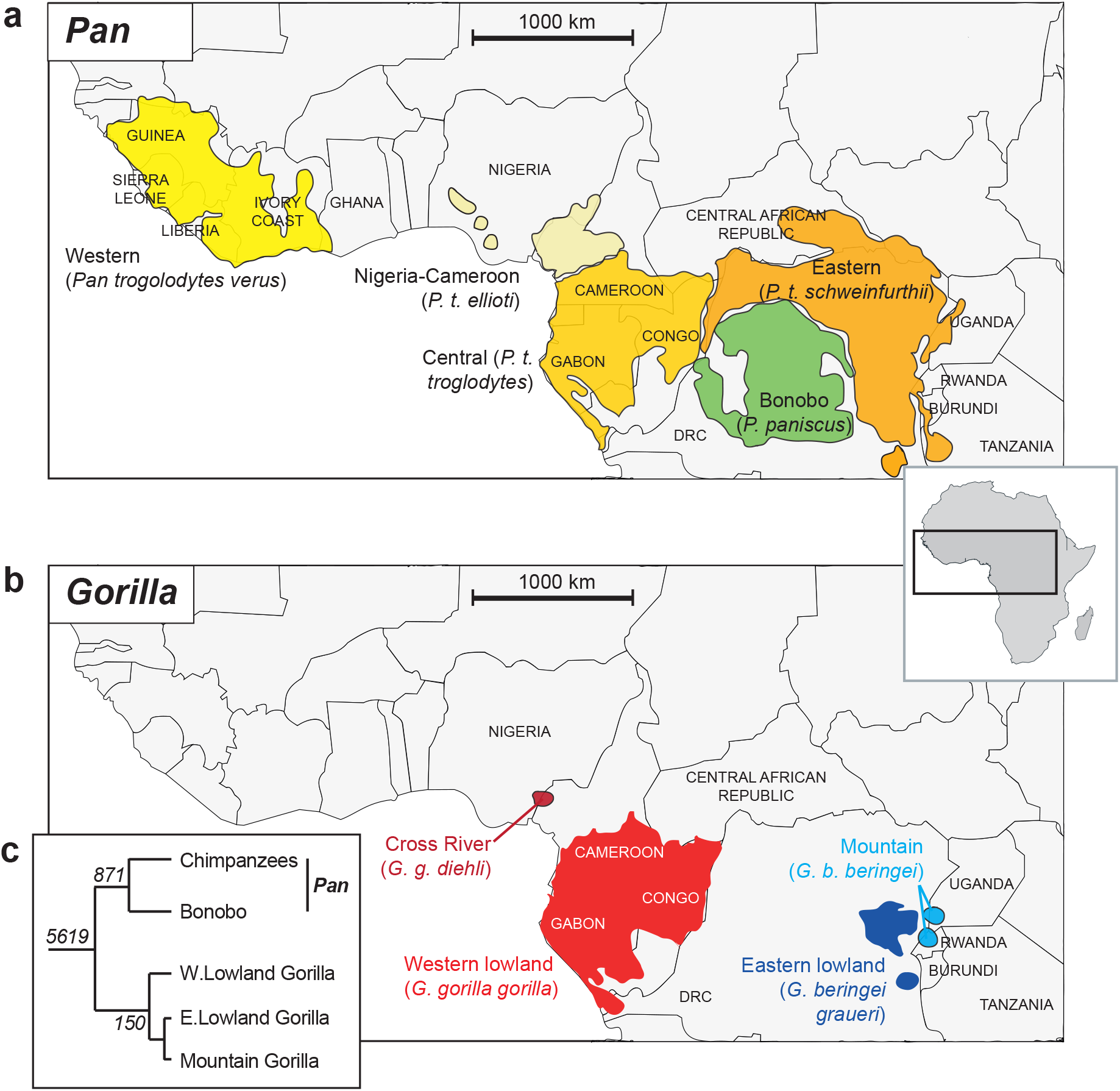
*Pan* and *Gorilla* species and sub-species distributions, and phylogenetic relationships. Distributions of a) *Pan,* and b) *Gorilla,* adapted from [60]. c) Phylogeny showing relationships between (sub)species (branch lengths not to scale), with classifications reflecting those used in this study. Italic numbers at nodes are split times in thousands of years, based on a mutation rate of 1 × 10^-9^ per bp per year [22], Map adapted from *Africa Just countries grayish.svg*, published on Wikimedia under a Creative Commons Attribution-Share Alike 4.0 International license. DRC: Democratic Republic of the Congo.

As a response, conservation efforts have seen an upsurge in the use of DNA testing to assess animal population parameters playing an important role in the implementation of effective wildlife management and preservation policies [8]. Measuring polymorphism at sets of autosomal short tandem repeats (aSTRs) via capillary electrophoresis (CE) has been an important tool in population genetic analysis [9]. Because they assort independently at meiosis, sets of unlinked aSTRs can also yield genotypes that provide individual identification within a species: this forms the basis of human forensic identification technologies [10], and can be applied in forensic casework involving animals, for example in poaching or illegal trade cases [11]. Such genotypes also have the potential to distinguish between species and subspecies when allelic spectra are suitably differentiated and characteristic.

Because of the high levels of sequence similarity among great-ape genomes [12], PCR primers for aSTR markers developed in humans are expected often to amplify their orthologs. Indeed, some STR multiplexes designed for human forensic analysis have been shown to have cross-species application for the analysis of orthologous loci in other great apes (e.g. [13, 14]). The underlying assumption is that amplicons generated at orthologous loci are generally commensurable across species [15]. However, this assumption is often incorrect; indeed, the presence of species-specific indels in flanking sequence together with different organisation and variability of STRs present difficulties with great-ape cross-species comparisons [15]. In translating multiplexes designed in humans to other species, there is also a practical problem of interpretation, since allele size ranges for different loci (labelled with the same fluorescent dye) were designed to be non-overlapping in humans, but may well overlap in non-human primates.

These issues arise because of the nature of capillary electrophoresis, which assesses polymorphism by measuring the length of PCR fragments and converting this to an assumed number of repeat units within each allele. An alternative approach is multiplex massively parallel sequencing (MPS), in which the sequences of STRs, rather than their lengths, are analysed. This obviates the problem of size-range overlap, since it is the sequence itself that identifies the locus, and also permits larger numbers of STRs to be simultaneously analysed than is possible with length-based CE genotyping. MPS-based analysis is now becoming established in human forensic genetics. For example, the ForenSeq DNA Signature Prep Kit (Verogen) [16, 17] includes multiple autosomal, X- and Y-chromosomal STRs, as well as autosomal SNPs for individual identification.

This study aims to assess how the human-designed ForenSeq multiplex system performs in amplifying and sequencing autosomal STRs in a set of chimpanzees, bonobos and gorillas, and to ask if the orthologous loci are both individually identifying and can robustly distinguish groups at the species and subspecies levels. Sequencing across subspecies and species may also reveal aspects of the mutation processes of these widely used STRs across several million years of primate evolution.

## Results

We assembled a set of DNA samples from 52 non-human great-ape individuals (14 chimpanzees, 4 bonobos, 16 western lowland gorillas, 6 eastern lowland gorillas, and 12 mountain gorillas) for sequencing. Both prior information on some sampled individuals and later deductions from our own data using the software ML-Relate [18] (Table S1) indicate that the sample set includes close relatives within (sub)species, including some parent-offspring, full-sib and half-sib pairs, though no mother-father-child trios. In describing the diversity of STR sequences and in considering identification at the individual and (sub)species levels, we retain all these individuals since they contribute new alleles to the dataset. When considering population structure, heterozygosity, inbreeding (*F*_is_), and forensic parameters we remove individuals such that there are no close relatives in the dataset, apart from in mountain gorillas. For this sub-species whole-genome sequencing [7] has shown chromosomes to be homozygous over >38% of their lengths; given this very high general relatedness, it seems reasonable to retain pairs suggested as half-sibs by ML-Relate.

### Amplification of orthologs of human loci in the multiplex

The ForenSeq^™^DNA Signature Prep Kit, with a human-based design, was used to amplify autosomal, X- and Y-STRs and autosomal SNP-containing loci in the set of 52 African great ape samples. Table S2 summarises amplification results across the entire set of 152 amplicons in the multiplex. Here, we focus on the 27 autosomal STRs (sequences given in Table S3), but also report the sequences of amplified X-STRs in Table S4. Not all samples are males, so Y-STR data are less extensive, and there is also a relatively high failure rate for amplifying orthologs of human loci. The amelogenin sex test loci [19] amplified in all individuals and gave results consistent with previously known sex (data not shown). We do not report sequence information for the human identity-informative SNP amplicons.

Of the 27 autosomal STRs targeted in the multiplex, two (D7S820 and D9S1122) failed to amplify in any individuals. D5S818 amplifies only in gorillas, but contains a low-diversity STR array with the structure (AGAT)_1-2_(AG)_9-13_, unlike the human ortholog which is a highly variable tetranucleotide repeat, (AGAT)_6-18_; we therefore do not consider it further here. Of the remaining 24 STRs, six (Figure 2) could be analysed only in particular species, likely due to inter-specific sequence differences affecting primer sites. A set of 18 STRs amplifies in all species, but with some missing data in particular individuals. Missingness could be due to null alleles arising from sequence variants affecting primer sites, or to poor sequence quality. Neglecting all STRs that show missing data leaves a ‘core’ set of thirteen STRs that were sequenced across all individuals; this set allows cross-species comparisons to be done.

**Figure 2:**
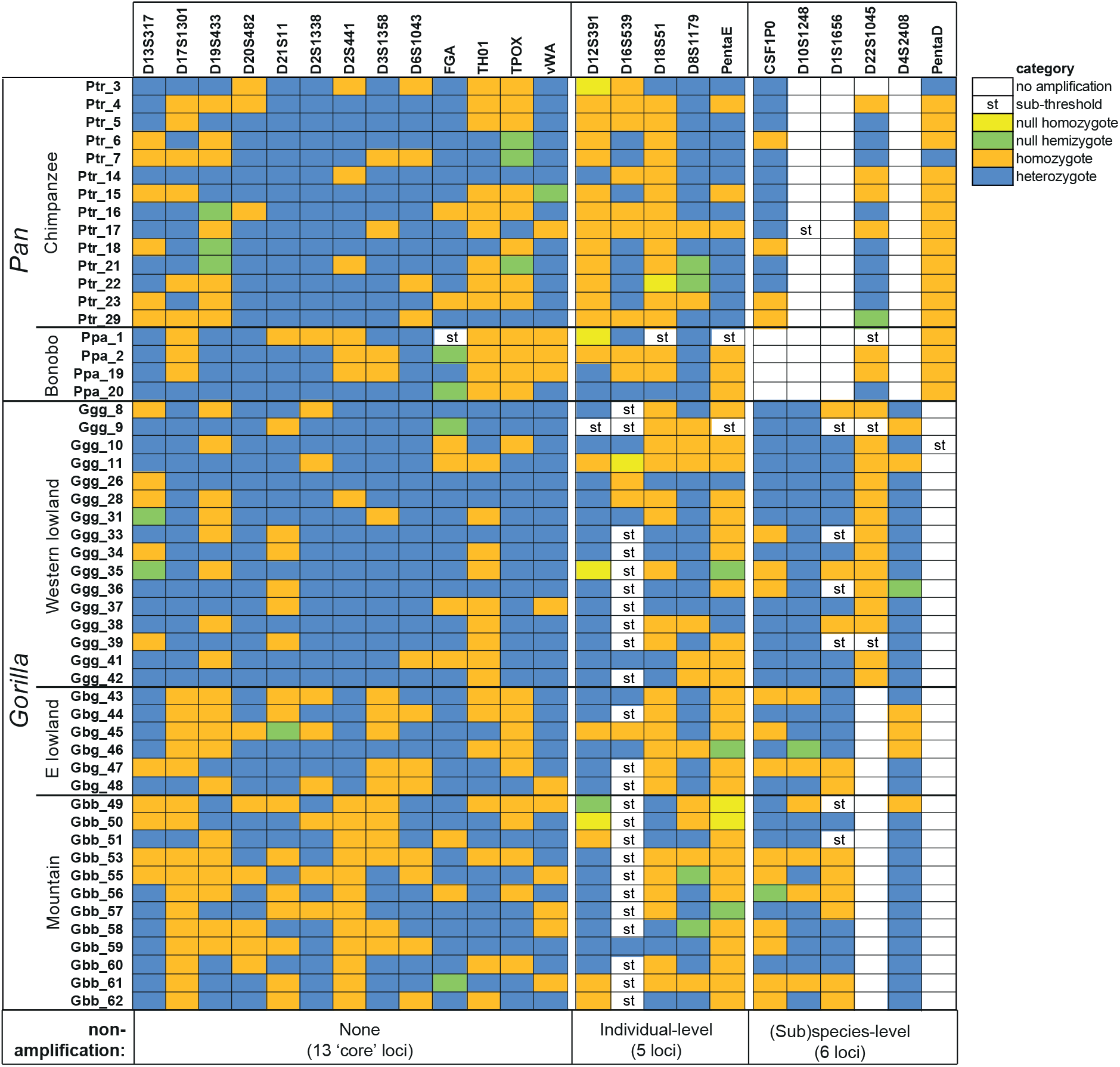
Summary of amplification behaviours of autosomal STRs across individuals. For each STR and each great-ape individual, amplification behaviour is summarised, as indicated in the key to the right. Distinction between categories is based on sequence read-depth analysis. STRs are organised into three groups reflecting the amplifiability and degree of data completeness as indicated below the figure.

### Sequence diversity in autosomal STRs

In humans, sequencing autosomal STRs increases allelic diversity [20, 21] by allowing variation within both the repeat array and flanking DNA to be observed (Figure 3a). This is also the case in the great apes studied here (Figure 3b-f; Table S5). Focusing on variation within the repeat array (since the lengths of flanking regions are not completely comparable between species) we see that STRs that show sequence variants are not well conserved across species. In humans, D12S391 shows by far the greatest increase in diversity due to repeat array sequence variation [20, 21], but this feature is not observed in the great apes studied here. D2S1338 shows the greatest degree of repeat array sequence variants across (sub)species.

**Figure 3:**
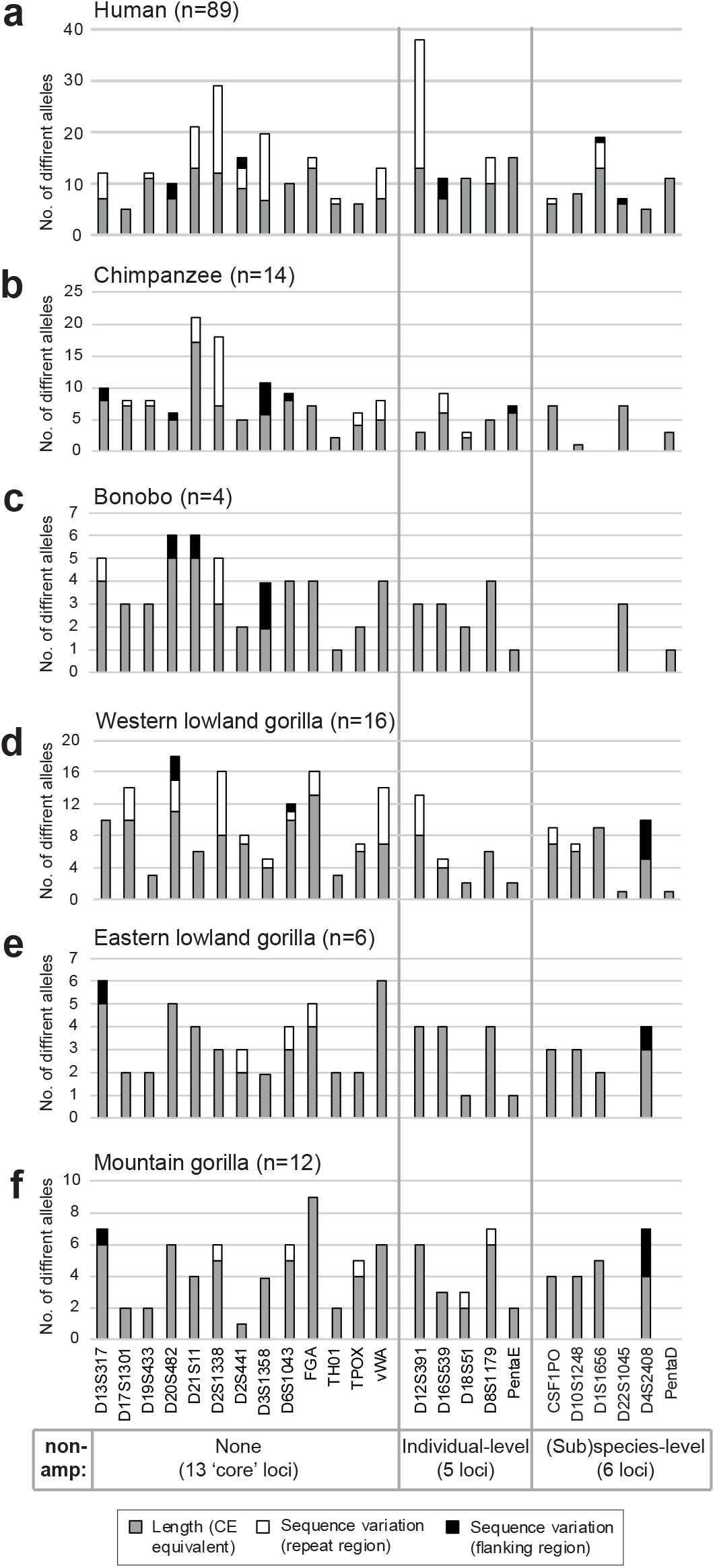
Counts of distinguishable alleles in each (sub)species by STR locus, and per-locus increment due to sequence variants. The observed numbers of length variants among individuals are shown as grey bars, and the number of additional alleles resulting from sequence variation within and flanking the repeat array are shown in white and black respectively. STRs are organised into three groups as in Figure 2, and shown below the figure. a) Human [21]; b) Chimpanzee; c) Bonobo; d) Western lowland gorilla; e) Eastern lowland gorilla; f) Mountain gorilla. Note that, although repeat array sequence variation is comparable across species, flanking sequence variation is not strictly comparable because the amount of sequence considered in different species varies somewhat.

### STR variant classes within and between (sub) species

To consider the sequence variation in the 18 cross-species amplifiable STRs in an evolutionary framework, we compared the *Pan* and *Gorilla* data to the predominant sequence structures of human orthologs (retrieved from STRBase.nist.gov and [20]), as well as to a single orangutan orthologous allele for all loci (where this could be identified), extracted from the orangutan *(Pongo abelii)* reference sequence (ponAbe3 assembly). Figure 4a summarises the STR structural categories observed; the range of allele structures for each locus is shown in a phylogenetic context in Figure 4b-h and Figure S1.

**Figure 4:**
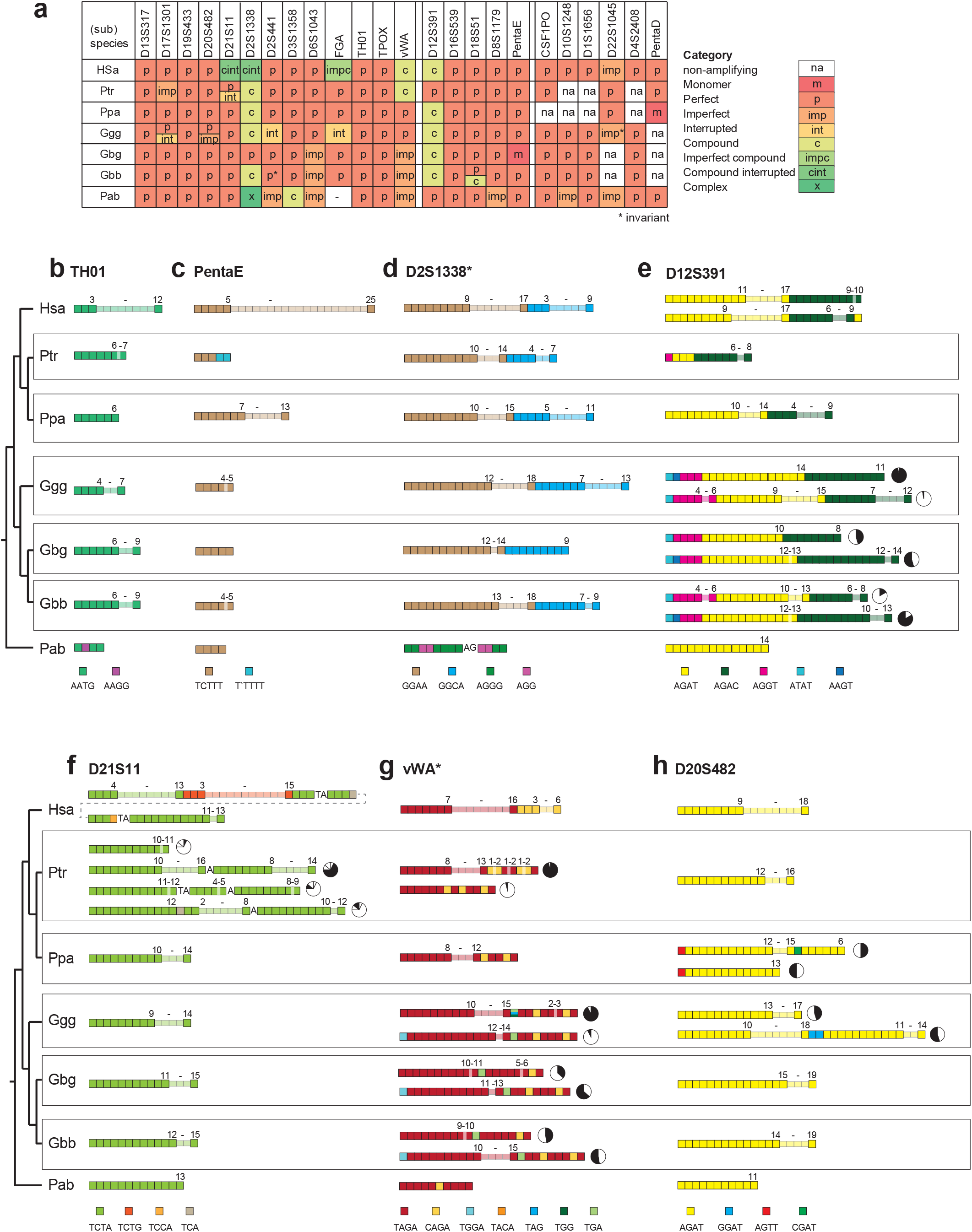
Summary of STR structures across (sub)species, and examples of inter- and intra-specific structural variation. a) For each (sub)species and each locus, the structural class of the STR is summarised as indicated in the key to the right. In cases where two classes are both present at high frequencies, the two classes are given as a split cell in the table. Human structures are taken from the predominant observed class listed at STRBase.nist.gov. Orthologous orangutan (Pab: *Pongo abelii)* alleles are based on the reference sequence. Hsa: *Homo sapiens;* Ptr: *Pan troglodytes*; Ppa: *P. paniscus*; Ggg: *Gorilla gorilla gorilla*; Gbg: G.*beringei graueri*; Gbb: G. *b. beringei*. b - h) Examples of variation across (sub)species, phylogenetically arranged, for seven STRs (see also Figure S1 for further examples). Human structures are from STRBase.nist.gov and [20]. In each case, tetra- or trinucleotide repeat motifs are indicated by boxes coloured according to the keys below. Ranges of repeat numbers within variable arrays are indicated. Where more than one structural class is observed within a *Pan* or *Gorilla* (sub)species, pie-charts indicate their proportions.

Several loci (including D13S317, D19S433, TH01, TPOX and D16S539) show evolutionary conservation across the great apes, with perfect repeat arrays of the same repeat unit across all (sub)species examined, and similar repeat ranges (Figure 4, Figure S1). We see no examples in which the major variable repeat unit differs in sequence between (sub)species, but among the remaining loci there is variation in structural types and little obvious relationship with the phylogeny, suggesting stochastic origins of mutations giving rise to diverse non-perfect repeat arrays. Repeat array length distributions are particularly well understood in humans because of very large sample sizes, whereas our great-ape sample sizes are small and may be highly unrepresentative. However, given this caveat, the number of repeats observed in all species fall within the range of human variation, with the exception of D13S317 (based on the lists given by STRBase.nist.gov and [20]).

Here, we summarise some features of structures for the 18 STRs that were amplifiable and sequenced across *Pan* and *Gorilla.* Many of these evolutionary comparisons appear to confirm anthropocentric ascertainment bias: for several STRs (in particular D6S1043, D18S51, D19S433, PentaE and TH01), recorded human allele repeat number ranges are much wider than those seen in our sample of great apes. In fact, across all 18 STRs, there is only one case, D13S317 in western lowland gorilla, where the observed non-human primate allele size range exceeds that seen in humans. While this may be influenced by the relatively large surveyed human sample sizes, it seems surprising given that, for example, western lowland gorillas have an effective population size more than twice that of humans [22], and it may suggest differences in mutational processes.

a. Some loci behave simply across the phylogeny with a lack of variant structures, and straightforward patterns of variation in perfect arrays. An example is TH01 (Figure 4b), which is a simple, perfect array of AATG repeats in humans, and the same across *Pan* and *Gorilla,* albeit with narrower repeat number ranges (and invariant in bonobos). The orangutan allele is very short and interrupted, and unlikely to be variable. Similarly simple behaviour is seen at PentaE (Figure 4c), D18S51, and D19S433 (Figure S1).
b. Two of the human loci, D2S1338 and D12S391, are compound in humans with two variable blocks of different repeat types. These features are conserved: D2S1338 (Figure 4d) shows similar structure and approximate array length ranges in humans, *Pan* and *Gorilla,* as a compound and polymorphic [GGAA]_n_[GGCA]_m_ STR. Surprisingly, the orangutan allele here comprises short arrays of different repeat units (AGGG and AGG). D12S391 (Figure 4e) shows variable arrays of AGAT and AGAC repeats, and in orangutan is a simple perfect array of just one of these repeat types, AGAT.
c. There is little evidence of novel repeat arrays arising and expanding in particular species. One exception is D21S11 (Figure 4f), which in all species shows one or more arrays of TCTA repeats, but in humans also includes a highly variable array of TCTG repeats that is not seen in any other species. The other example is at D12S391 (Figure 4e), where (as well as the AGAT and AGAC arrays mentioned above) an array of AGGT repeats is specific to *Gorilla,*and polymorphic in western lowland and mountain gorillas.
d. STR mutation processes are generally thought of as rapid compared to singlenucleotide changes in non-repetitive material, and (unless there has been recent gene flow) we might therefore expect little identity-by-descent in the features of repeat arrays over the several million years of primate evolution. However, this is not so, and the distribution of structures identical by descent appears to be non-uniform across the great apes. There are no examples of distinctive Pαn-specific derived features in any of the 18 STRs analysed. However, the picture is different in *Gorilla*. For D12S391 (Figure 4e), vWA (Figure 4g), D2S441, D16S539, FGA, and TPOX (Figure S1), all gorilla (sub)species studied carry more than one allele structure, and these are shared among western lowland, eastern lowland and mountain gorillas (which have an estimated divergence time of ~150 KYA; Figure 1c). Only one locus, D8S1179 (Figure S1), shows distinctive structural features restricted to the two eastern subspecies.
e. Considerations of STR array evolution based on human diversity and pedigree data have shown that interrupting a long perfect repeat array with a variant repeat or indel leads to a marked reduction of mutation rate [23] and consequent lower allelic diversity. However, in both *Pan* and *Gorilla* there are several allele structures featuring polymorphic arrays separated by interruptions (variant repeats, or insertions). In most of these cases, other variant structures in the same (sub)species are short and perfect, and these are shared across species suggesting they may be ancestral. This suggests that the long interrupted alleles might arise via a non-slippage-like process. Chimpanzee shows this phenomenon at D21S11 (Figure 4f) and D17S1301, while it is seen in *Gorilla* at D20S482 (Figure 4h), D13S317, D17S1301, and FGA (Figure S1).

### Within-(sub)species variability of multilocus STR genotypes

Within (sub)-species, all individuals (including related individuals; Table S1) are distinguishable by their STR genotypes, and this is true for both CE-equivalent and sequence-based allele designations.

After removing related individuals (Table S1) we assessed observed vs expected heterozygosity for the tested loci (Table S6); following Bonferroni correction, only one locus in one species (D16S539 in chimpanzee), shows a significant deviation from expectation. To ask if the tested loci reveal any evidence of inbreeding we estimated *F*_is_ (Table S7); following Bonferroni correction, significant positive *F*_is_ values are seen for three loci (D16S539, D19S433, TPOX) in chimpanzee, two (D13S317, D16S539) in western lowland gorilla, and one (D8S1179) in eastern lowland gorilla. As shown in Figure 2, all except one of these (TPOX) show evidence of null alleles or low read-depth in the relevant (sub)species, suggesting that the *F*_is_ results reflect technical issues rather than evidence of inbreeding. Forensic statistics derived from the data are given in Table S8, and Table 1 presents the combined random match probabilities (RMPs) in each (sub)species. The values obtained strongly reflect the sample sizes, which in turn influence the mean number of alleles observed per locus. RMPs are in all cases lower for MPS than CE allele designations, and in the range 10^-8^ to 10^-18^. Any comparison with human RMPs, where sample sizes and numbers of observed alleles are much larger, is not very meaningful. For example, the 24 loci analysable in western lowland gorillas give respective RMPs for CE- and MPS-based designations of 1.49 × 10^-27^ and 1.98 × 10^-30^ in a sample of 89 Saudi Arabian humans [21].

**Table 1:** Observed per genotype combined RMPs for different great ape (sub)species. See Table S8 for per-locus details.

| (sub) species | Mean no. chromosomes | No. loci | Mean no. alleles/locus |  | Combined RMP |  |
| --- | --- | --- | --- | --- | --- | --- |
|  |  |  | CE | MPS | CE | MPS |
| **Ppa** | 6.6 | 20 | 3.0 | 3.1 | 6.91E-08 | 5.15E-08 |
| **Ptr** | 20.9 | 22 | 5.8 | 7.0 | 7.78E-15 | 2.17E-15 |
| **Ggg** | 22.6 | 24 | 6.0 | 7.3 | 1.46E-17 | 1.61E-18 |
| **Gbg** | 10.8 | 22 | 3.0 | 3.2 | 6.21E-10 | 5.70E-10 |
| **Gbb** | 15.7 | 22 | 4.1 | 4.3 | 3.96E-12 | 8.53E-13 |

### Between-(sub)species variability of STR genotypes

To compare multilocus STR genotypes for the 13 ‘core’ loci across (sub)species, we carried out cluster analysis using STRUCTURE and DAPC, both for data at the full sequence level and for CE-equivalent (length-based). In STRUCTURE analysis of CE-equivalent data (Figure S2a), the best-supported value of *K* is 2, in which *Pan* and *Gorilla* form two clusters. DAPC analysis reveals three clusters, with *Gorilla* divided into clusters corresponding to western and eastern species (Figure S2b), reflecting the behaviour of this method in minimising differences within, while maximising differences between, populations. In STRUCTURE analysis of sequence-level data, *K* = 4 is best supported (data not shown), differentiating clusters corresponding to bonobo, chimpanzee, western gorilla and eastern gorilla (Figure 5a). DAPC analysis gives five clusters, separating out the two eastern gorilla subspecies (Figure 5b). Sequence-based analysis therefore performs better in distinguishing between (sub)species. Given the sharing of repeat motif variation across *Gorilla* (sub)species (Figure 4; Figure S1), it seems likely that the differences contributing to differentiation here reflect variation in the flanking sequences.

**Figure 5:**
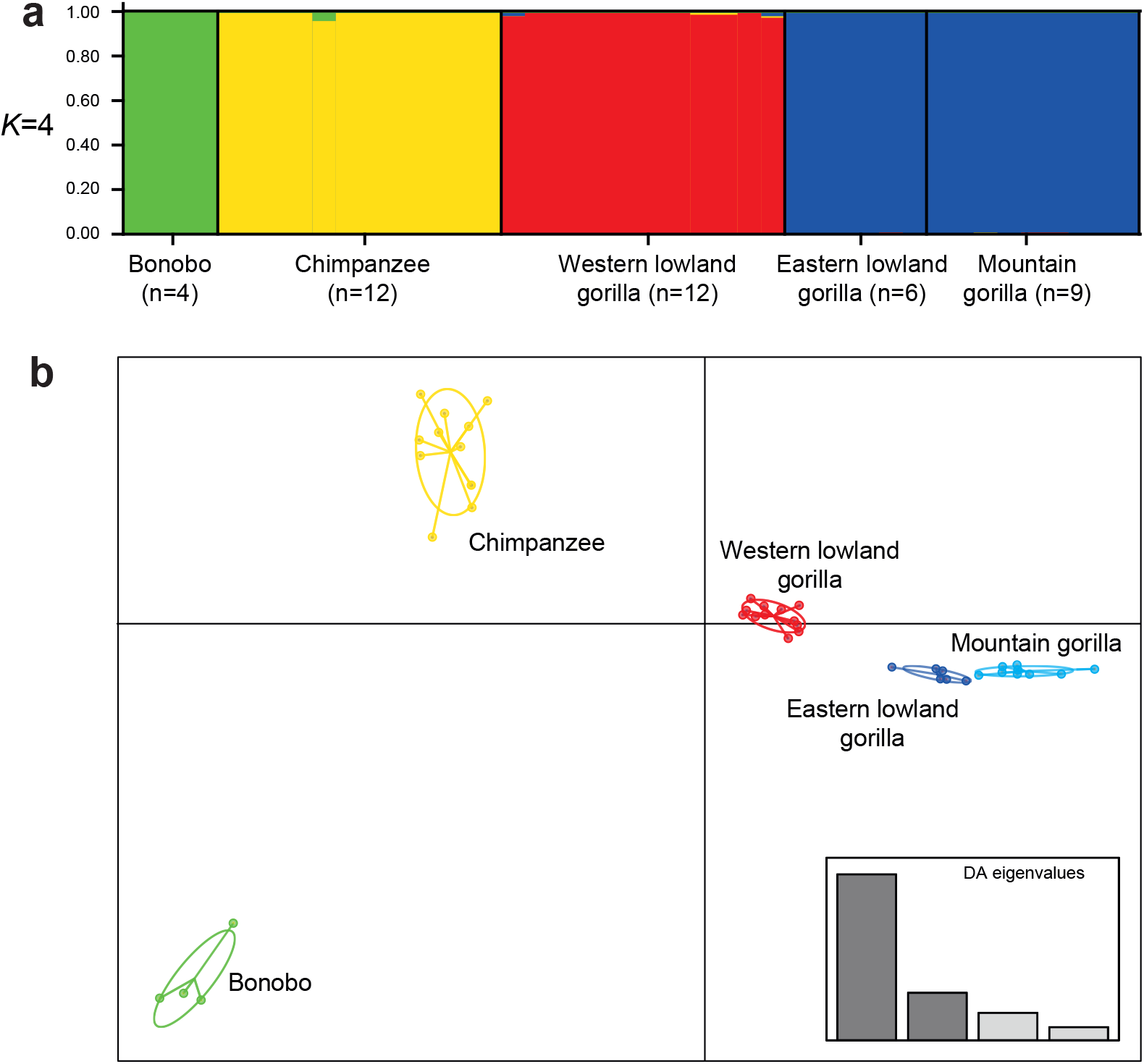
Cluster analysis based on sequence-based autosomal STR genotypes. a) Results based on STRUCTURE, for *K*=4; b) Results based on DAPC analysis. Full sequence information was used here (both array and flanking sequence data). An analysis based on CE-equivalent data is given in Figure S2. Related individuals are removed for this analysis (see Table S1).

## Discussion

Recent conservation initiatives have witnessed a considerable increase in the use of DNA testing for the implementation of effective wildlife conservation and management plans throughout the world. The current rate of biodiversity loss has prompted researchers to utilise markers that can be readily transferred between species to facilitate the study of taxa in which allelic diversity is poorly characterised [24, 25]. In this context, aSTRs have been a dominant source of neutral genetic markers for a variety of applications, including individual identification, assessment of population diversity and structure, and evolutionary studies [26]. Cross-species amplification depends on the presence of flanking sequences that, despite sometimes long divergence times, are conserved across organisms, and is directly related to the phylogenetic distance between the source and the target species [27, 28]. This has enabled the exploitation of common sets of PCR primers to type orthologous aSTR loci via capillary electrophoresis (CE) for the study of non-model organisms [15, 26, 29–32]. Following CE, PCR fragment lengths are converted into numbers of repeats at STR regions to produce individual genotypes. Recent studies, however, have identified several caveats to this approach, especially when it is used in cross-species analyses. Firstly, owing to convergent mutations, repetitive regions that are identical by state (i.e. have the same length) may not be identical by descent [33], therefore estimates of differentiation across species can be inaccurate. Secondly, CE fails to distinguish indels occurring within STR flanking sequences from changes in the structure of the repetitive regions, compromising the assessment of the organisation and variability of STRs [15]). As a result, the underlying assumption, under which orthologous STRs are commensurable across species, is often incorrect.

In recent years, the advent of MPS has obviated these problems by allowing researchers to investigate the structures of STR alleles, virtually in unlimited numbers. Consequently, MPS tolerates size homoplasy and the occurrence of overlapping ranges between loci that arise when homologous primers are used to genotype different species, as both STR and flanking sequences may not be invariant across species [15, 26, 29–32]. Because MPS does not rely on length discrimination, primer pairs can be strategically designed to target shorter fragments and increase multiplexing capability, thus making this technology particularly suitable for the analysis of highly degraded DNA found in non-invasive samples (e.g. faeces and hair [34]).

Here, we applied the human-designed ForenSeq kit to amplify and sequence human loci of forensic interest in 52 DNA samples from chimpanzees, bonobos, and gorillas, focusing on the results obtained for 27 autosomal STRs (aSTRs). As expected, given the low average sequence divergence between African great ape genomes (~1.3% between human and chimpanzee/bonobo [35]; ~1.75% between human and western lowland gorilla [36]), most of the aSTRs amplified successfully in most cases. Thirteen STRs could be genotyped in all individuals, and a further five showed only individual-level dropouts or sub-threshold amplification. The remaining nine either failed amplification altogether or failed in a particular species or genus. Failure to amplify is likely due to sequence divergence in primer-binding sites, though we have not investigated this since the ForenSeq kit’s primer sequences are proprietary and therefore not known exactly.

Our results show that MPS analysis of STR alleles can provide accurate individual and sub-species identification. In addition, STR structures show evidence of allele stability over long evolutionary times and reveal unexpectedly high levels of IBD across shared gorilla alleles (the only exception being D8S1179 in eastern gorilla subspecies, which reflects the short divergence time). Contrary to what has been reported in human pedigrees [23], we found that long interrupted alleles share a high degree of polymorphism across species, suggesting possible differences in mutation processes between species. In the future, increasing wholegenome sequence data at the population level and the application of genome-wide STR calling tools (e.g. HipSTR [37], LobSTR [38]) should illuminate these questions further.

Despite the advantages of MPS, the widespread adoption of high throughput sequencebased STR typing for wildlife conservation purposes is still hindered by high start-up costs (e.g. for equipment and reagents), labour-intensive sample preparation, and steep learning curves associated with MPS data analysis [39–41]. Additionally, the lack of well-established research facilities in biodiverse countries means that biological samples must be shipped to sites where sequencing can be performed [42]. Stringent international restrictions on the export of endangered species biological samples further contribute to increasing the cost and time of sequencing, *de facto* limiting the feasibility of DNA testing for wildlife conservation purposes [43, 44].

Nevertheless, recent technological advances have circumvented these issues by greatly reducing the cost for the acquisition of sequencing and laboratory equipment, with positive repercussions for the implementation of wildlife conservation genomics initiatives [39, 40]. In this regard, the commercialisation of portable nanopore sequencing devices by Oxford Nanopore Technologies Plc. promises to revolutionise the field of molecular ecology by permitting *in situ* analysis of DNA samples [42, 45–48]. The shift from a laboratory-centralised workflow to on-site DNA analysis overcomes the fundamental challenge of transporting biological material to a site where sequencing can be performed [42]. While only few studies to date have assessed the applicability of the ONT MinION device for sequencing forensic STRs [34, 49–51], recent findings suggest that STR panels can be compatible with ONT sequencing platforms [39], which opens up new opportunities in the field of wildlife forensics and conservation genetics.

## Conclusions

Our results indicate that MPS via a human-designed kit represents an effective method for the analysis of orthologous aSTR loci in non-human great ape species, and it provides reliable identification of individual and (sub)species. Comparison with standard length-based allele definitions shows higher observed allelic diversity and improved (sub)species discrimination.

## Methods

### DNA samples and data

DNA samples were from a variety of sources including laboratory collections, detailed in Table S1. For chimpanzees, subspecies definition was sometimes unclear, and where it was defined, sample sizes for individual subspecies were small: we therefore considered chimpanzees at the species level. By contrast, gorilla samples were better defined, with at least six individuals in each of three of the four known subspecies, and therefore gorillas were considered at this level. As a result, our comparison groups were five in number: chimpanzee - *Pan troglodytes* (n=14), bonobo - *P. paniscus* (n=4), western lowland gorilla - *Gorilla gorilla gorilla* (n=16), eastern lowland gorilla - *G. beringei graueri* (n=6) and mountain gorilla - *G. b. beringei* (n=12). To provide comparative information on the same set of loci in humans we used a published dataset based on analysis of the ForenSeq^™^DNA Signature Prep Kit in 89 unrelated Saudi Arabian human males [21], as well as information from STRBase.nist.gov and [20].

### Library preparation and sequencing

DNA samples were quantified using the Qubit^™^ Fluorometer with the Qubit^™^ dsDNA HS (High Sensitivity) Assay Kit for double-stranded DNA (dsDNA). Sequencing libraries were prepared with the human-based ForenSeq^™^DNA Signature Prep Kit according to the manufacturer’s recommendations (Verogen®, San Diego, CA, USA). Primer mix A was used to target 58 STRs (27 autosomal STRs, 7 X-STRs and 24 Y-STRs) and 94 identity-informative SNPs (iiSNPs) from 1 ng of template DNA. Steps for library preparation include amplifying, indexing, purifying, normalising and pooling, prior to sequencing on an Illumina MiSeq FGx, all of which were performed in accordance with the manufacturer’s recommended protocols.

### Sequence data analysis

Quality-checked FASTQ files were generated using Trimmomatic v.0.36 [52] for adapter sequence and poor-quality base trimming using the Linux terminal. The threshold for minimum read length was set at 50 bp.

Analysis of human data using the DNA Signature Kit is usually undertaken using the ForenSeq^™^ Universal Analysis Software (UAS), but for the non-human analysis done here the software FDSTools [53] was employed. This is laborious, but has the advantage that tailored anchor, flanking and repeat-array sequences can be designed, hence obviating the need for a reliable reference genome, which is still lacking for *Gorilla beringei* and most *Pan* sub-species. In order to develop library files for variant calling for *Pan* and *Gorilla,* trimmed bam files were visualised and aligned with the human reference (GRCh38/hg38) using the Integrative Genomic Viewer (IGV) [54] allowing the identification of suitable flanking sequences as anchors [53]. Considering the kit chemistry, which produces short and unreliable second reads, the 5’ anchor was set close to the 5’ end of the repeat array of each locus, so as to maximise the coverage for each marker. Flanking sequences to be added to the final version of the library file were obtained through repeated runs of FDSTools.

If there are null (non-amplifying) alleles at a given STR locus, these may exist in a heterozygous state, and it then becomes necessary to distinguish between such heterozygotes and true non-null homozygotes, in which two identical alleles are amplified. This was done using a sequence read-depth approach, normalised against known heterozygote calls, since a true homozygote’s read-depth should be equal to the sum of two heterozygous alleles (following a previous approach used for duplicated Y-STR alleles [55]).

### Population, forensic and statistical analysis

STRAF [56] was used to calculate forensic statistics, including genotype count (N), allele count based on sequence (N_all_), observed and expected heterozygosity (H_obS_ and H_exp_), polymorphism information content (PIC), match probability (PM), power of discrimination (PD), power of exclusion (PE), and typical paternity index (TPI).

Clustering of genetically similar individuals was investigated using both STRUCTURE [57], and discriminant analysis of principal components (DAPC). DAPC was conducted using the package adegenet (version 2.1-3) [58] implemented in R version 3.6.3 [59]. For DAPC, the function find.clusters() was used to determine the optimal cluster number without prior information, and the Bayesian information criterion (BIC) was used to identify abrupt changes in fit models for successive runs of increasing k-means clustering with *K*=1-8. The number of PCs to retain was cross-validated using the function xvalDapc() with 50 repetitions in order to avoid overfitting.

ML-Relate [18] was used to screen the sample set for closely related individuals within (sub)species. Based on this, together with some prior information (Table S1) some individuals were removed for some analyses, as described in the first paragraph of the Results section.

In considering STR repeat arrays across species, we consider four basic types: perfect (an uninterrupted array of a single repeat type, e.g. [GATA]_n_); interrupted (two or more arrays of the same repeat type interrupted by non-repeat material, e.g. [GATA]_n_NNNNN[GATA]_m_); imperfect (two or more arrays of the same repeat type interrupted by repeat-derived material, e.g. [GATA]_n_GACA[GATA]_m_ or [GATA]_n_GAT[GATA]_m_); compound (two or more variable arrays of different repeat types of the same length, e.g. [GATA]_n_[GACA]_m_). We also include two hybrid categories, compound interrupted (two or more variable arrays of different repeat types of the same length, interrupted by non-repeat material, e.g. [GATA]_n_NNNNN[GACA]_m_), and imperfect compound (two or more arrays of different repeat types of the same length, interrupted by repeat-derived material, e.g. [GATA]_n_AATA[GACA]_m_).

## List of abbreviations

aSTR: autosomal short tandem repeat
CE: capillary electrophoresis
DAPC: discriminant analysis of principal components
MPS: massively parallel sequencing
PCA: principal component analysis
PCR: polymerase chain reaction
SNP: single nucleotide polymorphism
STR: short tandem repeat
UAS: Universal Analysis Software

## Declarations

### Ethics approval and consent to participate

This research was approved by the University of Leicester’s Animal Welfare and Ethical Review Body (ref.: AWERB/2021/159), and all great ape samples were taken by qualified veterinarians.

## Consent for publication

Not applicable.

## Availability of data and materials

Most data generated or analysed during this study are included in this published article (and its supplementary information files). Additional raw data are available from the corresponding authors on reasonable request.

## Competing interests

The authors declare that they have no competing interests.

## Funding

EF was supported by a PhD studentship from the Natural Environment Research Council CENTA doctoral training programme (grant no. NE/L002493/1).

## Authors’ contributions

**EF:** Investigation, Methodology, Formal analysis, Visualisation. **JHW:** Conceptualisation, Supervision. **MAJ:** Conceptualisation, Supervision, Resources, Visualisation. **All authors:** Writing: original draft, reviewing and editing.

## Acknowledgements

We thank Yahya Khubrani and Tunde Huszar for assistance. We gratefully acknowledge colleagues who contributed DNA samples, particularly Chris Tyler-Smith and Yali Xue, and Lisa Gillespie (Twycross Zoo). We thank NUCLEUS Genomic Services at the University of Leicester for access to Illumina MiSeq sequencing. This research used the SPECTRE High Performance Computing Facility at the University of Leicester for data analysis.

## Notes

### Competing Interest Statement

The authors have declared no competing interest.

