## Supplementary Figures 1-2 for "Sequencing the orthologs of human autosomal forensic short tandem repeats provides individual- and species-level identification in African great apes"

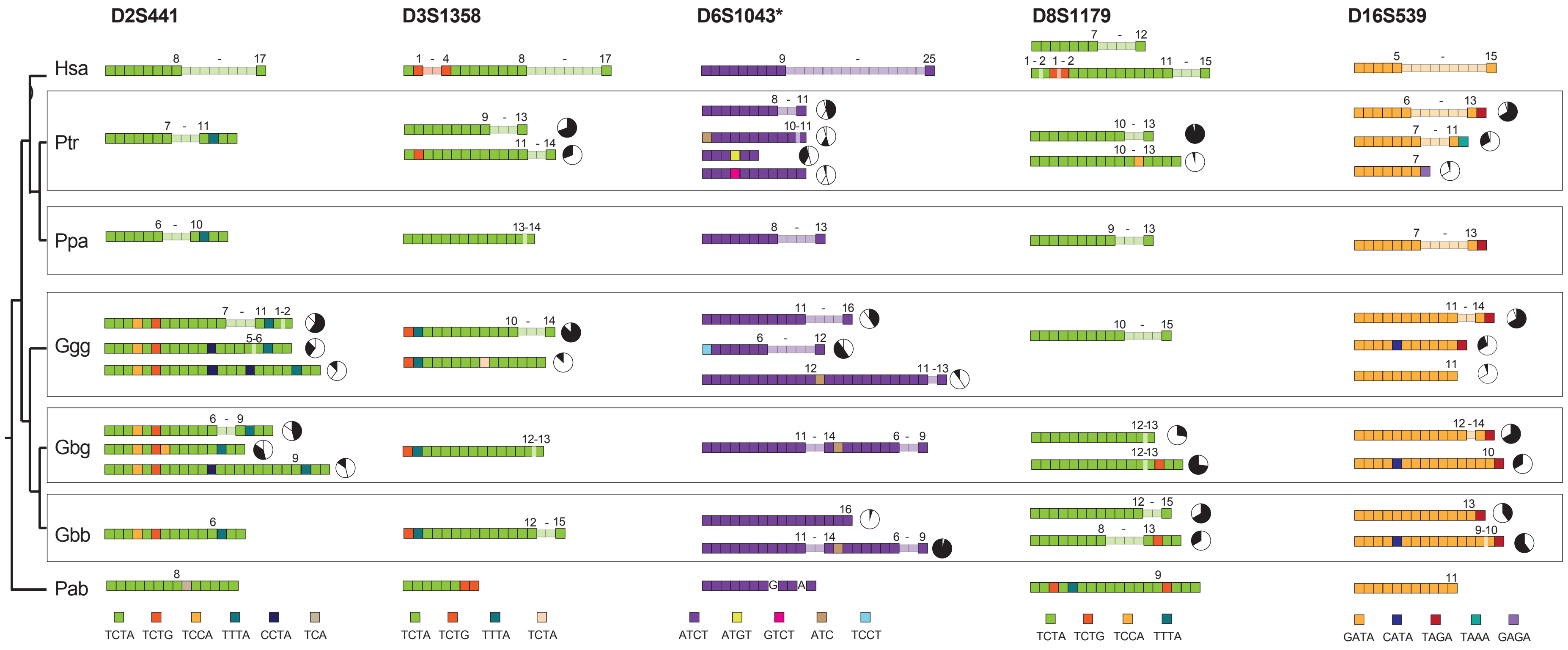


**Figure S1: Summary of inter- and intra-specific structural variation at 10 STRs (*continued on next page*).**

Schematic representation of variation across (sub)species, phylogenetically arranged, for ten STRs (see also Fig. 4). Human structures are from strbase.nist.gov and Gettings et al. (2016); for FGA, two highly divergent structures are shown. Orthologous orangutan (Pab: *Pongo abelii*) alleles are based on the reference sequence; none is observed for FGA. In each case, tetra- or tri-nucleotide repeat motifs are indicated by coloured boxes. Ranges of repeat numbers within variable arrays are indicated. Where more than one structural class is observed within *Pan* or *Gorilla*, pie-charts indicate their proportions. Hsa: *Homo sapiens*; Ptr: *Pan troglodytes*; Ppa: *P. paniscus*; Ggg: *Gorilla gorilla gorilla*; Gbg: *G. beringei graueri*; Gbb: *G. b. beringei*.

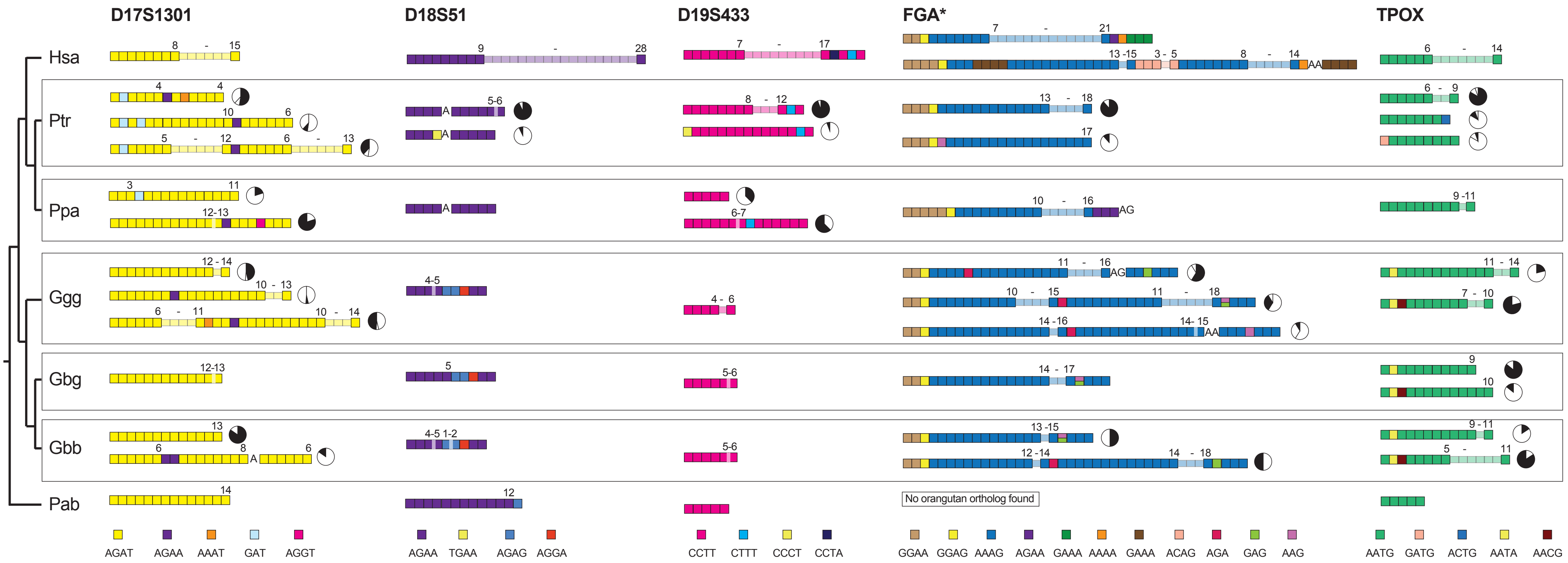


**Figure S1: Summary of inter- and intra-specific structural variation at 10 STRs (*continued from previous page*).**

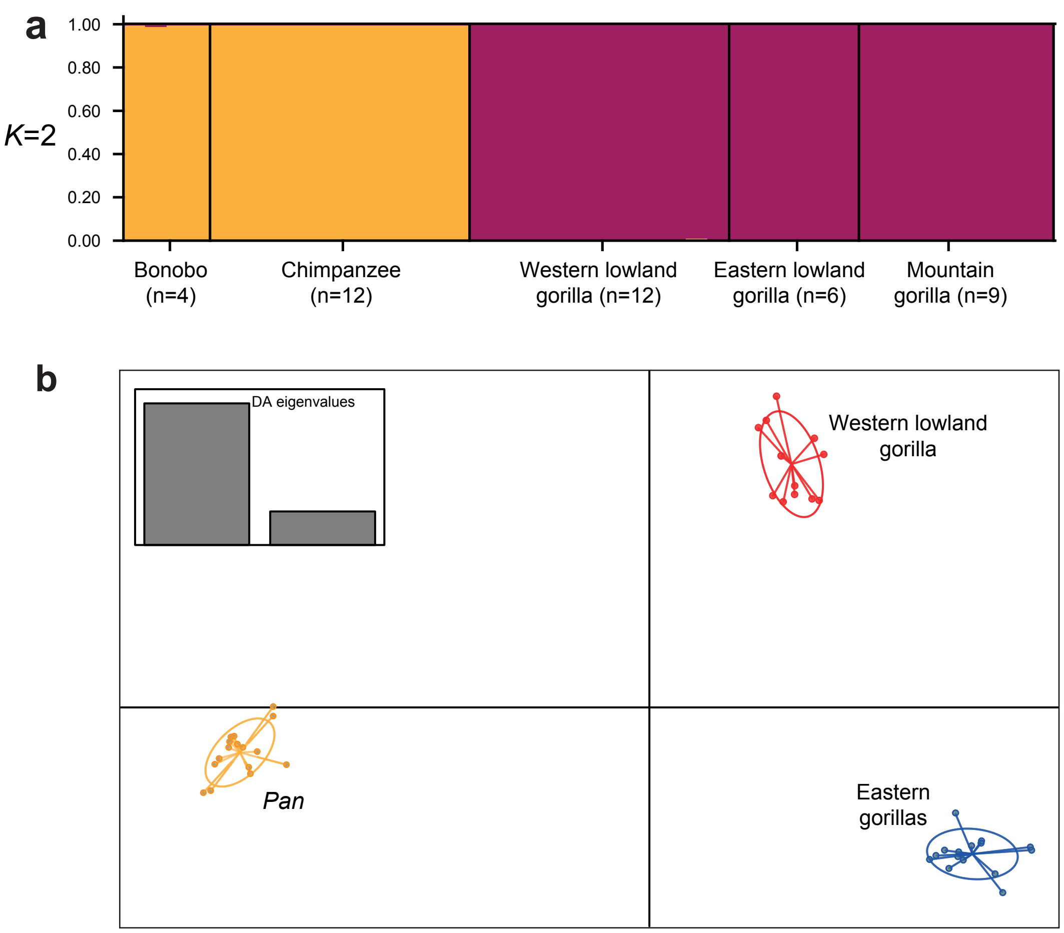


**Figure S2: Cluster analysis based on CE-equivalent autosomal STR genotypes.**

a) Results based on STRUCTURE, for *K*=2; b) Results based on DAPC analysis. Full information from MPS data was used here. An analysis based on sequence data (both array and flanking sequence variation) is shown in Figure 5. Related individuals are removed for this analysis (see Table S1).
